# Bioengineered human skeletal muscle with a Pax7+ satellite cell niche capable of functional regeneration

**DOI:** 10.1101/2020.05.04.076828

**Authors:** J.W. Fleming, A.J. Capel, R.P. Rimington, P. Wheeler, O.G. Davies, M.P. Lewis

**Affiliations:** School of Sports, Exercise and Health Sciences, Loughborough University, Loughborough, United Kingdom, LE11 3TU

**Keywords:** Skeletal Muscle, Regeneration, Tissue Engineering, Satellite Cell

## Abstract

Skeletal muscle (SkM) regenerates following injury, replacing damaged tissue with high fidelity. However, in serious injuries non-regenerative defects leave patients with loss of function, increased re-injury risk and often chronic pain. Progress in treating these non-regenerative defects has been slow, with advances only occurring where a comprehensive understanding of regeneration has been gained. Tissue engineering has allowed the development of bioengineered models of SkM which regenerate following injury to support research in regenerative physiology. To date however, no studies have utilised human myogenic precursor cells (hMPCs) to closely mimic human physiology due to difficulties generating sufficient cell numbers and the relatively low myogenic potential of hMPCs. Here we address problems associated with cell number and hMPC mitogenicity using magnetic association cell sorting (MACS), for the marker CD56, and media supplementation with fibroblast growth factor 2 (FGF-2) and B-27 supplement. Cell sorting allowed extended expansion of myogenic cells and supplementation was shown to improve myogenesis within engineered tissues and force generation at maturity. In addition, these engineered human SkM contained a Pax7+ niche and regenerated following Barium Chloride (BaCl_2_) injury. Following injury, reductions in function (87.5%) and myotube number (33.3%) were observed, followed by a proliferative phase with increased MyoD+ cells and a subsequent recovery of function and myotube number. An expansion of the Pax7+ cell population was observed across recovery suggesting an ability to generate Pax7+ cells within the tissue, similar to the self-renewal of satellite cells seen *in vivo.* This work outlines an engineered human SkM capable of functional regeneration following injury, built upon an open source system adding to the pre-clinical testing toolbox to improve the understanding of basic regenerative physiology.

## Introduction

Skeletal muscle possesses an innate and robust capacity to regenerate following injury, with most injuries regenerating the tissue to a state indistinguishable from that prior to injury (Järvinen *et al*, 2013). This regenerative capacity relies upon the presence of a resident stem cell population, satellite cells (SCs), which reside between the plasma membrane (sarcolemma) of muscle fibres and the encasing basement membrane (Mauro, 1961; Lepper *et al*, 2011; Hindi & Kumar, 2016). SCs are characterised by the unique position they occupy within the tissue, but also the expression of the stem cell transcription factor Pax7 (Seale *et al*, 2000). Following injury SCs are activated and proliferate readily (Dumont *et al*, 2015), producing committed myogenic precursor cells (MPCs) marked by the presence of MyoD expression (Tian *et al*, 2016; Zammit, 2017). MPCs then fuse together and with damaged muscle fibres, regenerating myofibres lost through injury (Bi *et al*, 2017; Quinn *et al*, 2017; Millay *et al*, 2013). In addition to a myogenic lineage, non-myogenic stem cells (fibro/adipogenic progenitors - FAPs) and immune cells support regeneration by modifying the extracellular matrix and coordinating repair and regeneration (Joe *et al*, 2010; Uezumi *et al*, 2010; Merly *et al*, 1999). The interactions between these additional cell types and MPCs has been shown to be vital in the regenerative process (Summan *et al*, 2006; Segawa *et al*, 2008; Wosczyna *et al*, 2019).

In severe muscle traumas such as volumetric muscle loss (VML), surgical trauma or partial muscles tears, the regenerative capacity of skeletal muscle can be overcome, leading to non-regenerative defects such as fibrosis (Mann *et al*, 2011; Olson & Soriano, 2009; Mueller *et al*, 2016; Fournier-Farley *et al*, 2016), interstitial adipose accumulation (Rose, 2002; Uezumi *et al*, 2010, 2011) and heterotopic bone formation (Belmont *et al*, 2012; Wheatley *et al*, 2015). In these circumstances individuals are left with complications such as reduced function, increased chance of reinjury and debilitating pain (Wangensteen *et al*, 2016; Dharm-Datta & McLenaghan, 2013; Freckleton & Pizzari, 2013). In animal models of severe traumas there has been limited success in manipulating the regenerative process to promote muscle regeneration and reduce non-regenerative defects. Although, where clear biochemical rationale exists, for example limiting the expansion of specific cell populations with small molecules, progress has been made (Fiore *et al*, 2016; Lemos *et al*, 2015; Agarwal *et al*, 2016). This limited success may be attributed to the lack of easily manipulated, high-throughput models of injury and regeneration, thus limiting understanding of the fundamental biology of muscle injury and regeneration/repair.

Due to the complex cell-cell interactions and three-dimensional (3D) environment necessary to accurately mimic skeletal muscle regeneration, models of muscle injury to date have been predominantly based around laboratory animals. However, animal models face limitations with low experimental throughput, complex genetic manipulations, complex pharmacological manipulation (compared to cell cultures), and ethical considerations, in addition to inherent biological variation from humans, and therefore there is a clear requirement to develop accurate and robust *ex vivo* models of pathophysiology (EU, 2010; Russell *et al*, 1959). Advances in tissue engineering have made it possible to create engineered skeletal muscle to understand complex physiological phenomenon. Engineered skeletal muscles from cell lines (Aguilar-Agon *et al*, 2019; Agrawal *et al*, 2017), primary laboratory animal MPCs (Morimoto *et al*, 2018; Huang *et al*, 2005; Vandenburgh *et al*, 1988), pluripotent stem cell (PSC) derived myocytes (Xu *et al*, 2019; Maffioletti *et al*, 2018; Rao *et al*, 2018) and primary human MPCs (Madden *et al*, 2015; Capel *et al*, 2019; Khodabukus *et al*, 2019; Bakooshli *et al*, 2019) have been demonstrated, but relatively few of these engineered skeletal muscles have been shown to possess a regenerative capacity following injury (Fleming *et al*, 2019; Juhas *et al*, 2014, 2018a; Tiburcy *et al*, 2019).

The regenerative processes of some engineered muscle models have shown clear correlations to that of *in vivo* muscle, and so the utilisation of these engineered tissues in studies of regeneration is a clear opportunity to increase our understanding of skeletal muscle regenerative physiology (Fleming *et al*, 2019; Juhas *et al*, 2018b; Tiburcy *et al*, 2019). Previous engineered models of regeneration have used the snake venom cardiotoxin (CTX) to induce a chemical muscle injury. CTX is widely used in animal models to produce a specific cellular model of muscle injury and has been shown to be effective in engineered tissues (Juhas *et al*, 2018b; Tiburcy *et al*, 2019). The model presented here takes a similar approach utilising Barium chloride (BaCl_2_), which is also widely used in laboratory animals as a specific myotoxin and produces a comparable injury type (Hardy *et al*, 2016). BaCl_2_ was chosen as an injurious stimulus due to previous *in vivo* publications, its high water solubility and ready availability with low regulatory restrictions, allowing easy and reproducible *in vitro* application (Mueller *et al*, 2016; Hardy *et al*, 2016). In addition, the mechanism of injury following BaCl_2_ treatment is a simple cellular injury specifically removing myotubes without reducing mononuclear cell number. Although chemical insults do not mimic all of the damage, specifically extracellular matrix destruction, caused by mechanical injuries usually seen *in vivo* these insults produce a specific and reproducible injury phenotype to ensure accurate model development (Fleming *et al*, 2019).

To ensure that data produced by these engineered models is as relevant as possible and that these models are exploited to their full potential, engineered muscles utilising primary human cells, with a regenerative capacity mimicking that of *in vivo* muscle, should be developed. To account for the heterogeneity of cells found within native muscle, primary tissue derived MPCs and not human iPSCs present the most biomimetic option for creating a representative model of human skeletal muscle regeneration.

Here we present a protocol to establish a high-throughput and robust, functional engineered human skeletal muscles from primary human MPCs. Utilising cell population sorting and media optimisation we present human engineered skeletal muscles which regenerate function and morphology completely following injury. These engineered muscles in addition to supporting regeneration also contain a self-renewing stem cell niche, presenting an opportunity to accurately study the biology of human skeletal muscle regeneration *ex vivo*.

## Materials and Methods

### Isolation and culture of hMPCs from skeletal muscle biopsies

Participants were recruited according to Loughborough University ethical and consent guidelines (Ethics no. R18-P098). Biopsies were collected by microbiopsy method from the *vastus lateralis* (Hayot *et al*, 2005). All collected tissue was minced finely, and connective tissue removed. Minced tissue was then plated out and cells isolated by explant culture (Martin *et al*, 2015; Capel *et al*, 2019). Once collected cells were expanded to passage 3 (p3) in gelatin coated (0.2% v/v) culture flasks. At p3 cells were sorted for the presence of the myogenic cell surface marker CD56 (Illa *et al*, 1992), using a MidiMACS™ system (Miltenyi Biotech, DE). Further expansion of the separate populations was then undertaken, CD56+ cells in Corning® Matrigel® basement membrane matrix coated (1mg/mL, Fisher Scientific, UK) flasks and CD56-cells in gelatin solution (Sigma Aldrich, UK) coated flasks. At p5 cells were cryopreserved or further expanded and used between p7 and p9. Throughout explant and expansion cells were maintained in growth media (GM – 79% High glucose Dulbecco’s Modified Eagle’s Medium (DMEM, Sigma, UK), 20% Fetal Bovine Serum (FBS, PanBiotech, UK) and 1% Penicillin/Streptomycin (P/S, Fisher, UK)). For the culture of minced tissue 1% Amphotericin B (Sigma, UK) was added to standard GM.

### Generation of tissue engineered muscles

Engineered muscles were made as described previously (Capel *et al*, 2019; Fleming *et al*, 2019). Briefly, 65% v/v acidified type I rat tail collagen (2.035mg/mL, First link, UK) and 10% v/v of 10X minimal essential medium (MEM, Sigma) were mixed and neutralised. This was followed by the addition of 20% v/v Matrigel® and 5% v/v GM containing hMPCs at a final density of 4×10^6^ cells/mL and in a ratio of 9:1 CD56+:CD56- unless otherwise stated. The final solution was transferred to pre-sterilised biocompatible polylactic acid (PLA) 3D printed 50μL inserts (Rimington *et al*, 2017) to set for 10-15 minutes at 37^°^C. All moulds used in this manuscript are freely available to download at the following URL: https://figshare.com/projects/3D_Printed_Tissue_Engineering_Scaffolds/36494. Engineered skeletal muscles were maintained in GM with 5ng/mL FGF-2 (Peprotech, US) for 4 days, changed every 48 hours. Following 4 days media was changed to differentiation media (DM – 97% DMEM, 2% horse serum and 1% P/S) supplemented with Gibco™ B-27™ Supplement (50X, 1:50, Sigma) for a further 10 days.

### Barium Chloride injury and regeneration

Once engineered muscles had reached maturity (14 days), as defined above, they were exposed to chemical injury by BaCl_2_. Prior to inducing injury, fresh DM was added to all conditions. Precisely 50μL/mL of 12% w/v BaCl_2_ solution was then added to the medium for injury culture conditions, followed by a 6 hour incubation to induce injury. Following injury, cultures were washed once with phosphate buffered saline (PBS) to remove residual BaCl_2_ containing media. Control (no injury) and 0-hour (0 Hrs) time points were collected at the end of injury incubation. Additional time points at 2-, 4-, 9- and 14-days post injury were collected for all measures to examine the regenerative response across time. For the first 4 days of regeneration engineered muscles were maintained in GM with fibroblast growth factor 2 (FGF2 aka bFGF), and the remaining 10 days DM with B-27.

### Tissue fixation, sectioning and staining

Engineered muscles were fixed in 3.75% formaldehyde solution overnight at 4^°^C, and then stored in PBS. Prior to cryosectioning, engineered muscles were stored in 20% sucrose solution w/v for 24 hours at 4^°^C to reduce water content and then were frozen under isopentane in liquid nitrogen. Sections were then prepared using standard cryotomy methadology. Cross sections for MyHC staining were prepared at 12μm, whilst Pax7 and MyoD staining used 4μm sections. Longitudinal sections were prepared at 10μm.

Sections were incubated overnight at 4^°^C with primary antibodies and 1 hour with secondary antibodies (Fisher, Goat anti-mouse/rabbit 488/647, 1:500). For Pax7 (deposited to the DSHB by Kawakami, A., US, 1:125) and MyoD (Santa Cruz Biotechnology, US, sc-377460, 1:200) antigen rescue at 75^°^C in citrate buffer (pH 6) for 20 minutes was performed before addition of primary antibody. For Laminin (Abcam, ab11575,1:200) and MyHC (deposited to the DSHB by Fischman, D.A., MF-20, 1:200) antigen rescue was not required. DAPI (Fisher, 1:1000) was used to stain nuclei.

Images were collected on a Leica DM2500 microscope using Leica Application Suite X software. Fiji 1.52e (Schindelin *et al*, 2012) was used for image analysis, and an in house macro performed automated myotube and nuclei analysis. Pax7 and MyoD positive nuclei analysis was performed manually. Five random images per repeat, per measure were taken and analysed to generate the presented data.

### RNA extraction and Real Time-Polymerase Chain Reaction (RT-PCR)

Engineered muscles were snap frozen upon collection and TRIReagent® extraction was augmented by mechanical disruption of constructs in a TissueLyser II (Qiagen, UK) for 5 minutes at 20Hz. Following disruption RNA extraction was carried using chloroform extraction, according to manufacturer’s instructions (TRIReagent®, Sigma). RNA concentration and purity were obtained by UV-Vis spectroscopy (Nanodrop™ 2000, Fisher).

All primers (Supp Table 1) were validated for 5ng of RNA per 10μL RT-PCR reaction. RT-PCR amplifications were carried out using Power SYBR Green RNA-to-CT 1 step kit (Qiagen, UK) on a 384 well ViiA Real-Time PCR System (Applied Bio-systems, Life Technologies, ThermoFisher, US), and analysed using ViiA 7RUO Software. RT-PCR procedure was: 50°C, 10 minutes (for cDNA synthesis), 95°C, 5 minutes (reverse transcriptase inactivation), followed by 40 cycles of 95°C, 10 seconds (denaturation), 60°C, 30 seconds (annealing/extension). Melt analysis was then carried out using standard ViiA protocol. Relative gene expressions were calculated using the comparative CT (ΔΔCT) method giving normalised expression ratios (Schmittgen & Livak, 2008). RPIIβ was the designated housekeeping gene in all RT-PCR assays and sample controls for each primer set were included on every plate.

### Measurement of engineered muscle function

Electric field stimulation was used in order to assess the functional capacity (force generation) of tissue engineered constructs. Constructs were washed twice in PBS, and one end of the construct removed from the supporting mould pin. The free end of the construct was then attached to the force transducer (403A Aurora force transducer, Aurora Scientific, CA) using the eyelet present in the construct. The construct was positioned to ensure its length was equal to that before removal from the pin and covered (3mL) with Krebs-Ringer-HEPES buffer solution (KRH; 10mM HEPES, 138 mM NaCl, 4.7mM KCl, 1.25mM CaCl_2_, 1.25mM MgSO_4_, 5mM Glucose, 0.05% Bovine Serum Albumin in dH_2_0, Sigma, UK). Aluminium wire electrodes, separated by 10mm, were positioned parallel either side of the construct to allow for electric field stimulation. Impulses were generated using LabVIEW software (National Instruments, UK) connected to a custom-built amplifier. Maximal twitch force was determined using a single 3.6V/mm, 1ms impulse and maximal tetanic force was measured using a 1 second pulse train at 100Hz at 3.6V/mm, generated using LabVIEW 2012 software (National Instruments). Twitch and tetanus data were derived from 3 contractions per construct, and a minimum of 2 constructs per time point per biological repeat. Data was acquired using a Powerlab system (ver. 8/35) and associated software (Labchart 8, AD Instruments, UK).

### Experimental repeats

For all injury experiments (Fig 4 and 5), 5 repeats across 3 donors were performed with each repeat yielding a minimum of 3 engineered muscles per analysis type. For cell composition and media supplementation (Fig 1 and 2) 2 repeats across 2 donors were performed with 3 engineered muscles per analysis technique. A total of 5 donors were used for the entirety of the experimental work.

**Figure 1:**
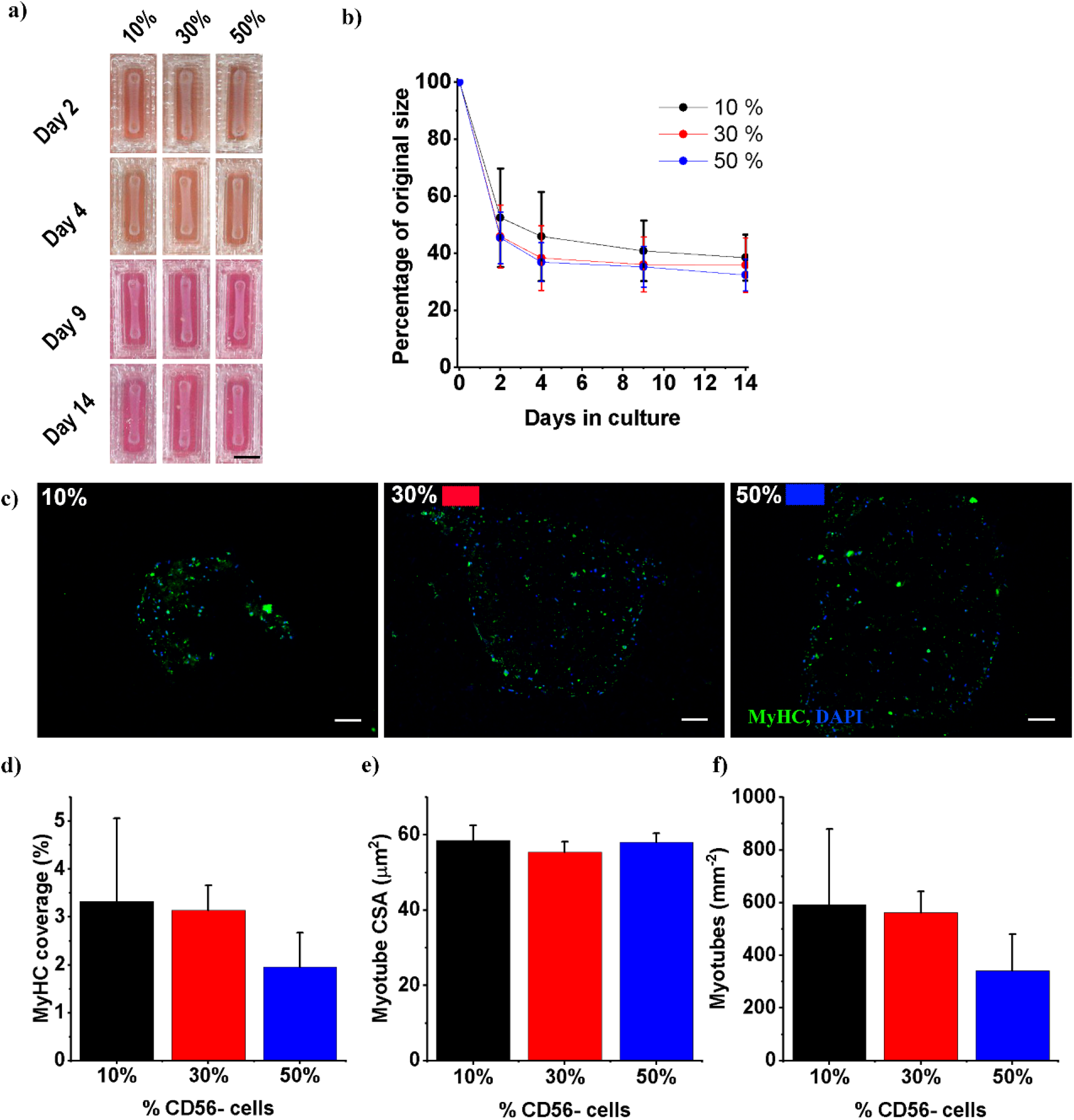
Remixing of CD56 sorted populations leads to robust engineered muscles. Throughout percentages refer to the percentage of CD56-cells remixed to create the human cell population. Black in all graphs denotes 10%, red 30% and blue 50% CD56− **(a)** Photographs of engineered muscles across time showing deformation. Scale bar – 5mm **(b)** Deformation over time **(c)** Representative micrographs stained for MyHC - green and Nuclei (DAPI) – blue. Scale bar 100μm. **(d-f)** Graphs displaying; MyHC percentage coverage, myotube cross sectional area (CSA) and myotubes per mm^2^. All graphs display mean ± S.D.

**Figure 2:**
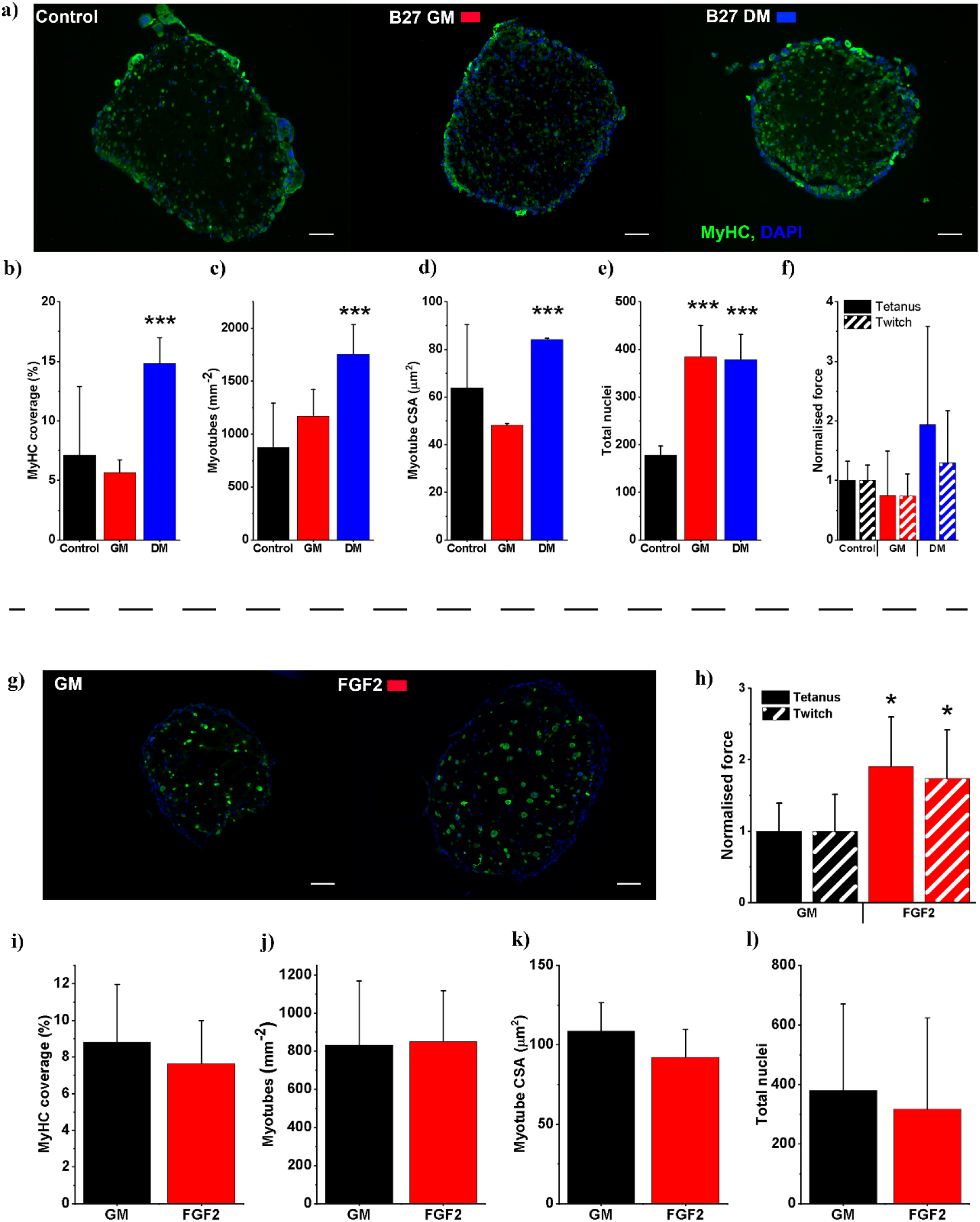
Media supplementation of remixed engineered muscles increases morphological maturity and increases functional capacity. **(a)** Representative micrographs stained for myosin heavy chain (MyHC) - green and Nuclei (DAPI) – blue. Scale bar 100μm. **(b-f)** Graphs displaying; Percentage MyHC coverage, myotubes per mm^2^, myotube cross sectional area (CSA), nuclei per section and normalised force (normalised to control) respectively. Mean tetanus force at control 72.2μN, twitch force 35.8μN. All graphs display mean ± S.D. **(g)** Representative micrographs stained for myosin heavy chain (MyHC) - green and Nuclei (DAPI) – blue. Scale bar 100μm. **(h)** Normalised force (normalisd to GM condition) mean tetanic force for GM condition 14.3μN, twitch 6.2μN. **(i-l)** Graphs displaying; Percentage MyHC coverage, myotubes per mm^2^, myotube cross sectional area (CSA), nuclei per section respectively. All graphs display mean ± S.D. Statistical significance is denoted as * p≤0.05, ***p≤0.001.

### Statistical analysis

Statistical analysis was undertaken in IBM SPSS 23. Data was subjected to tests of normality (Shapiro-Wilk) and homogeneity of variance (Levene’s Test). Where parametric assumptions were met, an ANOVA test was used to identify significant interactions. Where significant interactions were observed, Bonferroni post-hoc analyses were used to analyse differences between specific time-points or groups. Non-parametric Kruskal-Wallis analysis was undertaken where data violated parametric assumptions. Mann-Whitney (U) tests were then used, with a Bonferonni correction, to identify the differences between groups. Comparisons across time were made between control and the time point of interested and quoted p values refer to this comparison. All data is reported as mean ± standard deviation (SD). Significance was assumed at p≤0.05.

## Results

### Remixing CD56+ and CD56− cell populations produce robust tissue engineered muscles

The sorting of non-myogenic and myogenic populations from human explant cultures allows extended culture periods within the myogenic populations without a significant loss of desmin positivity, a marker of myogenic potential (Supp Fig 1b). However, the use of only myogenic cells in collagen/Matrigel® hydrogels produces engineered muscles of highly variable quality due to the apparent inability of these cells to reproducibly deform the hydrogel matrix (Supp Fig1c/d). However, due to the high proportion of myogenic cells in the CD56+ fraction (referred hereafter to as CD56+) constructs these engineered muscles, when successful, produce significantly more myotubes than unsorted equivalents (Supp Fig1 e-i).

To exploit the high myogenic potential of CD56+ cells, a dose remixing experiment was undertaken to identify the lowest proportion of CD56− cells required to reproducibly deform Collagen/Matrigel® constructs. A CD56− cell fraction was remixed with the CD56+ fraction at various ratios (10% 9:1 CD56+:CD56−, 30% 7:3 CD56+:CD56− and 50% 1:1 CD56+:CD56−) and hydrogel deformation and morphological appearance of the tissue examined. All ratios produced engineered muscles which deformed robustly, without any significant difference between conditions (Fig 1a/b). Clear trends in morphological appearance were present across conditions with 10% CD56− constructs displaying the highest number of myotubes per square mm and the largest percentage of constructs occupied by myotubes (Fig 1c,d/f). A small decrease between 10% and 30% CD56− and a much larger step between 30% and 50% CD56− engineered muscles was observed indicating that increasing the CD56− fraction reduced myogenic potential (Fig 1d/f). As the proportion of CD56− cells increased these measures of myotube formation were reduced, although this trend was not significant. Myotube cross-sectional areas (CSA) remained unaffected by the proportion of CD56− cell included in constructs (Fig 1e). As 10% CD56− remixing produced robust deformation and allowed the maximum inclusion of the CD56+ myogenic fraction this remixing ratio was carried forward for all future experiments.

### Media supplementation increases morphological and functional markers of muscle maturity

Engineered muscles (CD56−:CD56+, 10:90) were supplemented either in the growth phase (Day 0-4) or the differentiation phase (Day 4-14) of culture with 2% B-27 supplement. Supplementation with B-27 increased nuclei number approximately 2-fold irrespective of the phase in which it was added (p=0.001, p=0.004, Fig2e). However only supplementation in the differentiation phase of culture lead to increases in total MyHC coverage, myotube number and myotube CSA (p<0.001, p<0.001, p=0.004, Fig2 a-d). No significant changes in force generation were observed in any B-27 supplementation conditions, however supplementation in the differentiation phase of development did lead to a mean increase in tetanic force of 94%.

Supplementation of GM with FGF2 at 5ng/mL followed by 10 days in B-27 supplemented DM was compared to unsupplemented GM followed by B-27 supplemented DM. FGF2 supplementation did not change any morphological measure significantly (Fig 2 g, i-l). However, FGF2 addition did increase force generation significantly for both tetanus (1.9-fold, p=0.017) and twitch (1.74-fold, p=0.044). This data allowed the selection of FGF2 supplemented GM and B-27 supplemented DM as suitable medias for the culture of 10% CD56− engineered human skeletal muscles.

### Human engineered muscles display laminin organisation and Pax7+ nuclei

To identify how similar engineered skeletal muscle was in its matrix and cellular organisation to somatic muscle, features of *in vivo* muscle morphology were examined. We specifically examined the basement membrane organisation and the presence or absence of Pax7+ nuclei. Laminin staining of cross sections showed clear and distinct concentrations of laminin staining surrounding virtually all myotubes within engineered muscles, closely reminiscent of the basement membrane organisation of *in vivo* muscle (Fig 3b). Longitudinal analysis showed the presence of Pax7+ nuclei between a laminin rich area of matrix and the associated myotube (Fig 3a). This organisation was similar in organisation to the satellite cell niche of *in vivo* muscle and suggested that these Pax7+ cells may be functional and act as a myogenic precursor population in response to injury.

**Figure 3:**
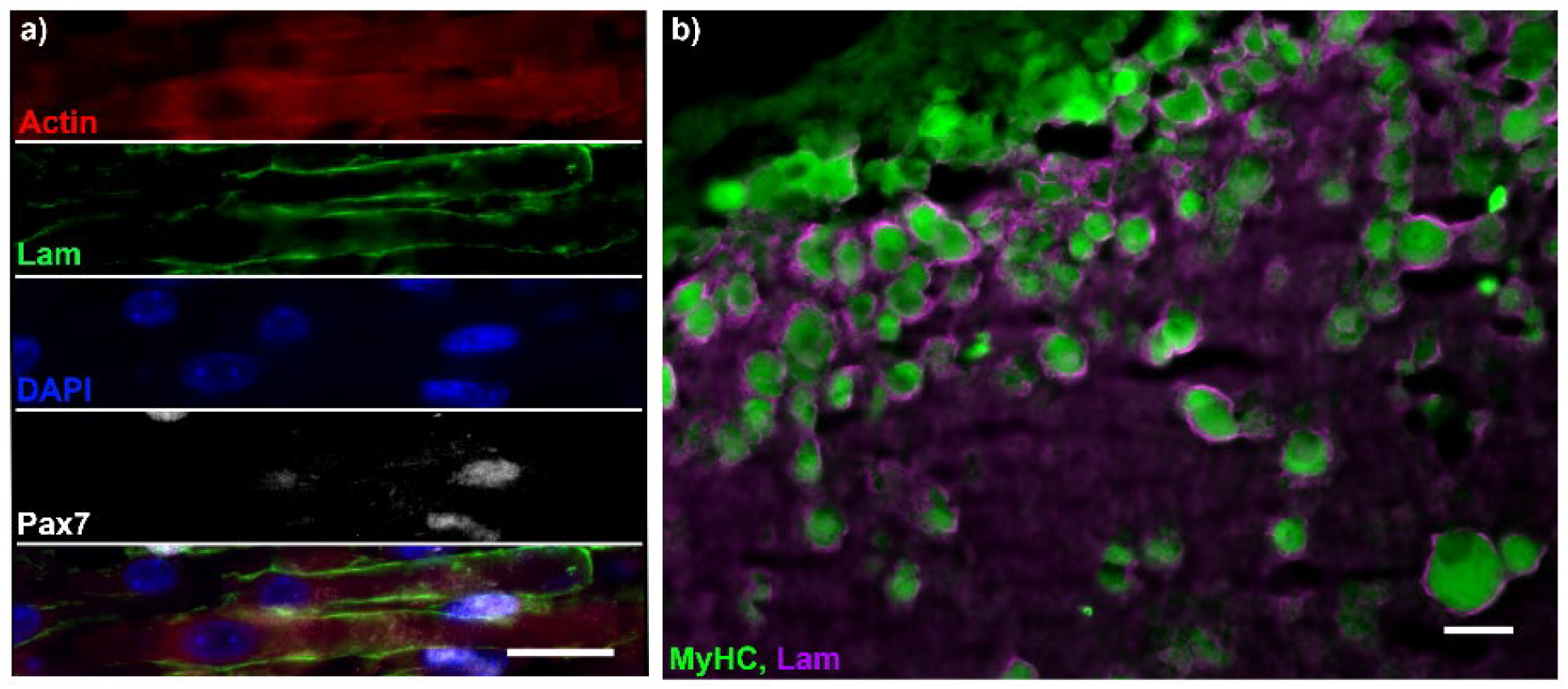
Human engineered muscles display laminin organisation and Pax7+ nuclei. **(a)** Micrograph of longitudinal cryosection of engineered muscle. Stained for Actin (Red), Laminin (Green), DNA (DAPI, Blue), and Pax7 (White) **(b)** Cross section of engineered muscle imaged on the extreme periphery of the section. MyhC (Green) and Laminin (Magenta). All scale bars represent 25μm

### Engineered skeletal muscles are capable of functional and morphological regeneration following injury

To examine the regenerative capacity, and so the function of the Pax7 niche, engineered muscles were exposed to an injurious stimulus in the form of BaCl_2_ exposure. Injury caused an initial reduction of 27.5% in MyHC positive coverage (p=0.006) and this reduction in coverage became more pronounced up to 4 days post injury (54.7%, p<0.001, Fig 4a/d). This reduction was caused predominantly by a loss of myotubes (myotubes per square mm), from 972 mm^−2^ pre-injury to 789 mm^−2^ immediately post injury (p=0.005, Fig4 a/d) further reducing to 638 mm^−2^ (p=0.001) 4 days post injury. This reduction was also accompanied by moderate atrophy of myotubes, with CSA reducing 9% following injury, although not significantly. This atrophy increased to a maximum of 29% (38.9μm^2^, p<0.001) at 4 days post injury. Across all measures of myotube size and density the largest reduction was seen between control and immediately post injury (6hr BaCl_2_ incubation), however these measures continued to decline across the first 4 days of regeneration. At 9 days post injury MyHC coverage had recovered to 85% of uninjured levels and was no longer significantly reduced. In addition, myotubes per square mm had returned to 94% of uninjured controls, suggesting that myotube CSA was still depressed. Indeed, myotube CSA at day 9 post injury was significantly reduced, with an average CSA of 115μm^2^ compared to 133μm^2^ at control (p=0.05, Fig 4a/d). Following the full 14 days of regeneration all measures of myotube density and size had returned to control levels showing complete morphological regeneration of the tissue.

**Figure 4:**
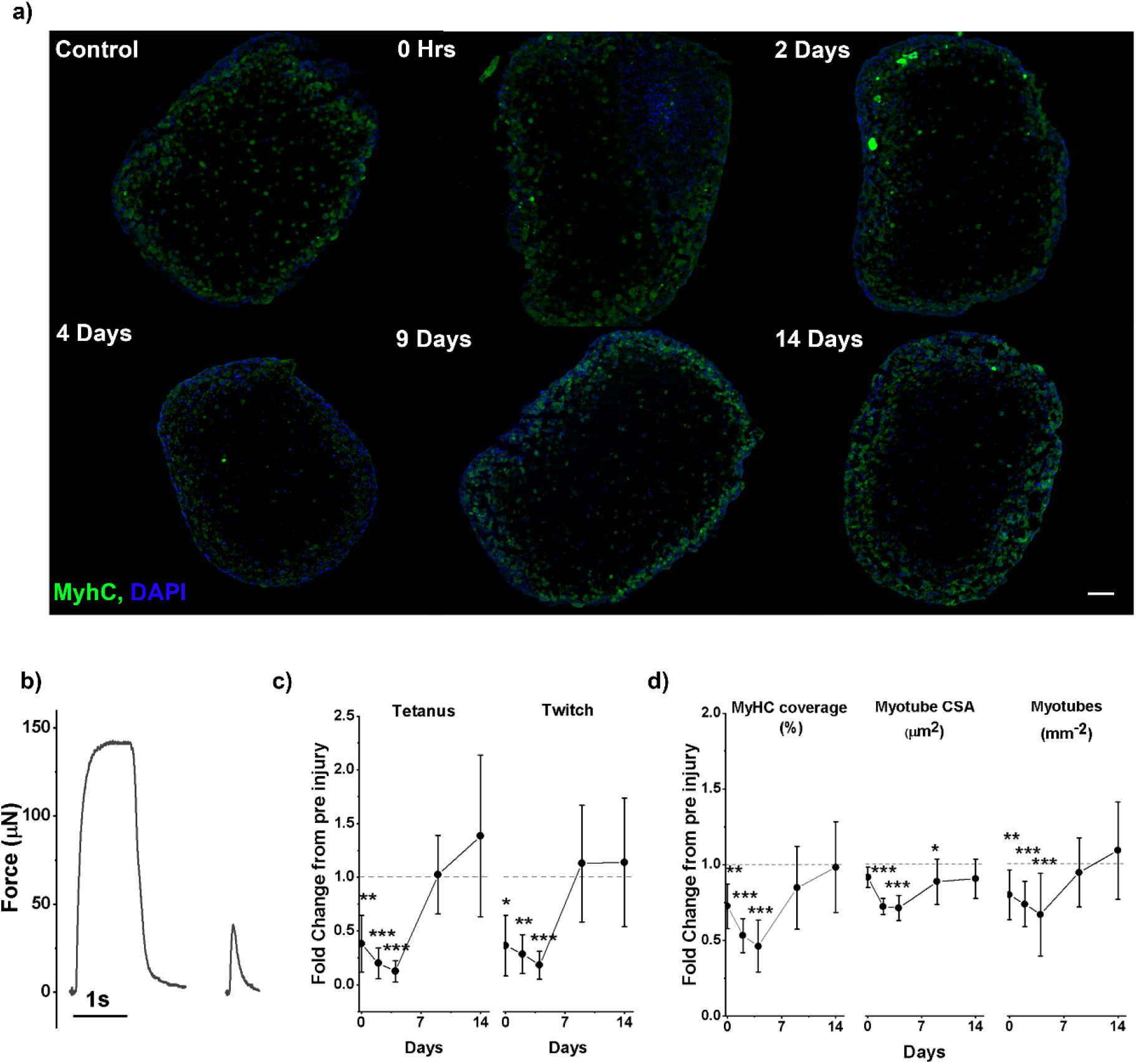
Human engineered muscles regenerate functionally and morphologically following injury. **(a)** Representative micrographs of engineered muscle cross sections. Stained for pan-MyHC (Green) and Nuclei (DAPI, Blue). Scale bar represents 100 μm. **(b)** Representative force traces used for Tetanus and Twitch force measurements. **(c)** Normalised force measurements across recovery. Means at control; Tetanus - 101.6 μN, Twtich - 36.6μN. **(d)** Normalised morphological measures across recovery. Means at comtrol;Precentage MyHC coverage - 13.0%, Myotube cross sectional area (CSA) – 132.6μm^2^, Myotubes per mm^2^ - 971.6. **(b-d)** All graphs display control normalised means ± S.D. Dashed line represents level at control, normalised to 1, on all graphs. Statistical significance is denoted as * p≤0.05, **p≤0.01 and ***p≤0.001.

With morphological change an accompanying reduction in functional output is expected. Immediately following injury both tetanic and twitch force were reduced by an average of 62% compared to control (p=0.003, p=0.024, Fig 4c). Both measures of function remained significantly depressed at 2-days and 4-days post-injury when compared to control. At day 9 post-injury both twitch and tetanic force were recovered to control levels and remained comparable to control at the end of regeneration at 14 days post injury (Fig 4c).

### Dynamics of myogenic and non-myogenic cell populations during regeneration

Nuclei per square mm was recorded to ensure that engineered muscles remained viable throughout recovery, and no significant variation in this measure was observed (Fig 5c). However, an increase of 15% was observed 2 days post injury (p>0.05), although this was resolved by 4 days post injury. To examine if different populations of cells within the overall population were expanding in relation to nuclear markers of the myogenic lineage; Pax7 and MyoD, were stained for and expressed as a percentage of total nuclei. MyoD, which marks proliferative myoblasts committed to the myogenic lineage, showed no change immediately following injury. However, a significant increase from 26.4% at control to 38.6% was observed after 2 days of regeneration (p=0.005, Fig 5a). This expanded MyoD population was completely collapsed following a further 2 days, at 4 days post injury, with the percentage of MyoD positive nuclei returning to 26.0%. Through the remainder of regeneration, the percentage of MyoD positive nuclei remained comparable to control. Pax7 a marker of satellite cells *in vivo* was found to be initially rare, making up only 0.42% of total nuclei at control, and no significant changes in Pax7 positive nuclei percentage was observed in the first 4 days following injury. However, following 9 days of regeneration Pax7+ nuclei comprised 2.4% of total nuclei a significant increase (p=0.002, Fig 5b) which persisted until 14 days post injury where Pax7 positive nuclei comprised 2.1% of total nuclei (p=0.004, Fig 5b).

**Figure 5:**
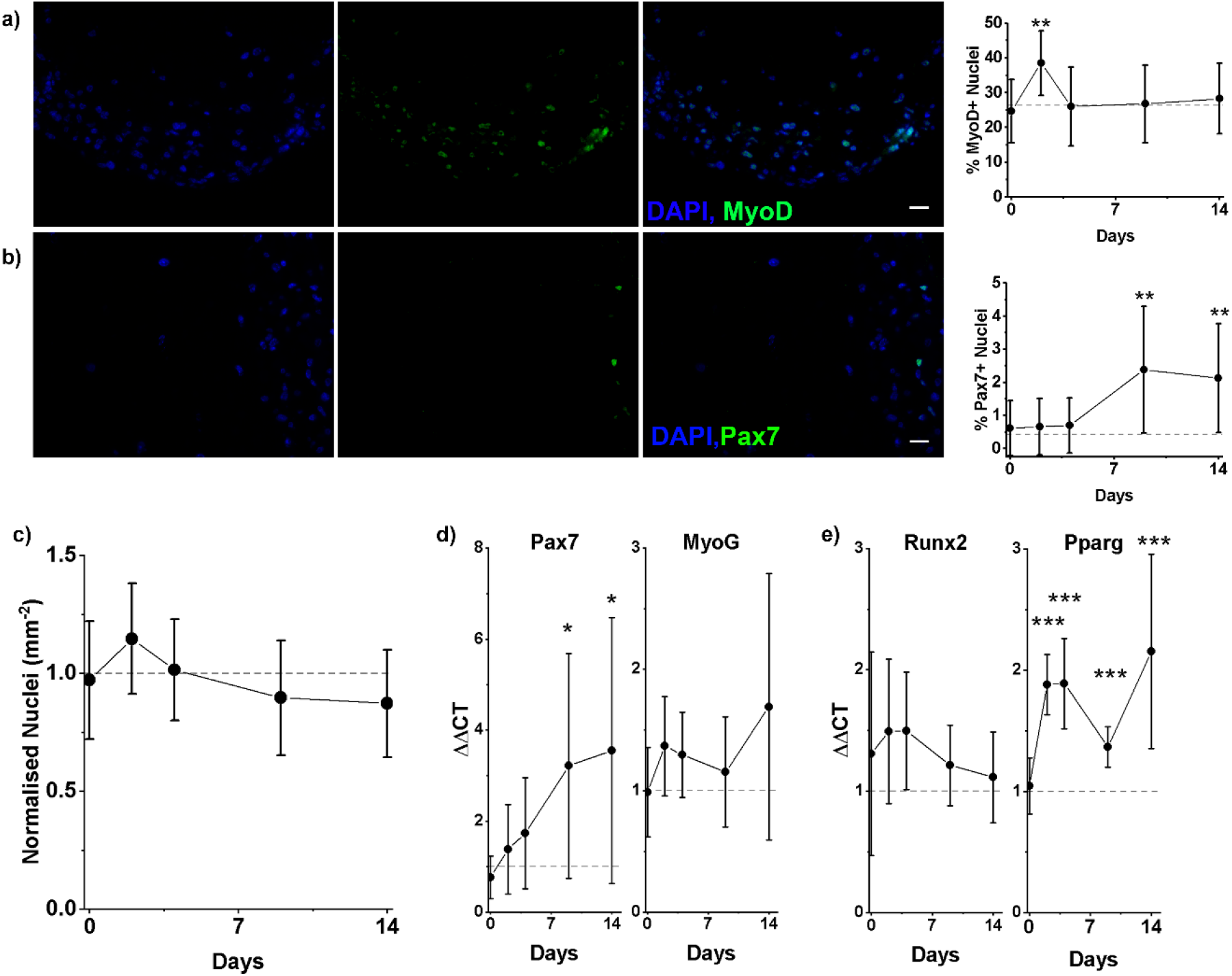
Dynamics of cell populations following injury. **(a)** Representative images showing staining left to right, Nuclei (DAPI, Blue), MyoD (Green) and overlay image. Scale bar represents 25μm. Graph displays percentage of MyoD+ nuclei across recovery. **(b)** Representative images showing staining left to right, Nuclei (DAPI, Blue), Pax7 (Green) and overlay image. Scale bar represents 25μm. Graph displays percentage of Pax7+ nuclei across recovery. **(c)** Normalised nuclei per mm^2^ across recovery, mean value at control-1471mm^−2^ **(d)** RT-PCR data displayed as ΔΔCT values for myogenic genes Pax7 and MyoG across recovery. **(e)** RT-PCR data displayed as ΔΔCT values for non-myogenic developmental genes Runx2 and Pparg across recovery. **(a-e)** All graphs display means ± S.D. Dashed line represents level at control on all graphs. Statistical significance is denoted as * p≤0.05, **p≤0.01 and ***p≤0.001.

Examination of Pax7 expression by RT-PCR showed a similar trend as Pax7 staining. Pax7 mRNA expression increased throughout recovery with 3.2 and 3.5-fold increases at 9- and 14-days post injury being significant (p=0.02, p=0.015). Although the 3-fold changes observed by RT-PCR are smaller than the approximate 6-fold change observed in staining the trends appear to be consistent, with increased Pax7 positive nuclei appearing through the final 10 days of regeneration (Fig 5d). Myogenin (MyoG) expression was also analysed, as a later myogenic marker than MyoD it would be expected to follow a similar trend but temporally delayed. The trend of MyoG expression is broadly similar to MyoD positive nuclei with a rise following 2 days of regeneration and partially resolving at day 9 post injury. An increase at 14 days post injury is observed although this is not statistically significant (Fig 5d).

The non-myogenic transcription factors Runx2 and Pparg were also examined to illuminate the dynamics of potentially non-myogenic cells which *in vivo* can drive the formation of non-regenerative defects such as increased interstitial fat and heterotopic bone. Runx2 mRNA showed no statistically significant deviation from control levels, although an increase was observed at all time points across the first 9 days of regeneration (Fig 5e). Pparg mRNA was significantly increased throughout regeneration, with the exception of immediately post injury. At 2- and 4-days post injury Pparg mRNA expression was elevated 1.88 and 1.89-fold respectively (p<0.001,p<0.001, Fig 5e). This expression was reduced to 1.37-fold at 9 days post injury (p<0.001, Fig5e) before elevating again to 2.16-fold (p<0.001, Fig 5e) after 14 days of regeneration. The upregulation of Pparg suggests a level of non-myogenic repair occurring in response to injury.

## Discussion

Here we have demonstrated by the exploitation of well-established tissue engineering approaches to develop a protocol which allows the generation of engineered human muscle containing a functional regenerative niche. The tissues generated are functional, contain the mixed cell populations represented in muscle, and demonstrate renewal and even expansion of the Pax7 niche through recovery. Although previous work has shown that engineered muscles can regenerate following injury (Juhas *et al*, 2014, 2018a; Fleming *et al*, 2019; Tiburcy *et al*, 2019), no previous work has demonstrated this with human engineered muscles. The injury sustained following BaCl_2_ treatment is robust, leading to significant loss of myotubes and function. As with *in vivo* skeletal muscle wound healing and previous primary tissue engineered muscles, injury is followed by a period (in this model 4 days) of impaired function and reduced myotube number (Hardy *et al*, 2016; Juhas *et al*, 2018a; Tiburcy *et al*, 2019). This contrasts with our previous cell line based 3D-models, which isolates the response of committed myogenic cells, and shows a very rapid recovery of function with no prolonged post-injury period (Fleming *et al*, 2019). Following this initial period without myotube formation a complete recovery of myotube number and function was observed in line with similar injury types in animal models and previous engineered tissues (Hardy *et al*, 2016; Fleming *et al*, 2019; Juhas *et al*, 2014; Tiburcy *et al*, 2019). This demonstrates the regenerative capacity of human skeletal muscle *ex vivo* and positions this model as a useful tool in examining the underlying biology of skeletal muscle regeneration and repair.

Initially we demonstrate that the remixing of CD56− and CD56+ cells is required to create robust engineered muscles in line with previous work (Brady *et al*, 2008; Bakooshli *et al*, 2019). This protocol demonstrates that only a very low proportion of non-myogenic cells are required to drive hydrogel deformation, allowing a high percentage of myogenic cells to be exploited (Fig 1). MidiMACS sorting is unlikely to be absolutely efficient in producing a population of homogeneous cells on the basis of CD56 expression, however, the desmin positivity values in excess of 90% observed here suggest a highly efficient sorting yield. In addition to increasing myogenic potential, sorting and remixing reduces donor to donor variation in desmin positivity - a common observation with hMPCs - reducing some of the donor variability and likely increasing experimental power. Finally, sorting allows separate expansion of myogenic and non-myogenic populations and therefore extends the usability of hMPCs. hMPCs unsorted rarely retain sufficient desmin positivity for engineered tissue after 6 passages, and therefore a standard microbiopsy sample would yield approximately 100×50μL engineered muscles. However, sorting cells allows further expansion (up to at least 9 passages, which could potentially be extended) and a projected 20-fold increase in MPC yield allowing 2000 engineered muscles to be generated per biopsy sample. Taken together these advantages suggest clear rational for exploiting CD56 sorting for hMPC tissue engineering and solves one of the key limitations of using hMPCs.

Remixing of separate populations produces a method to increase cell yield and reproducibility of engineered human skeletal muscle by allowing the expansion of competing populations in isolation. However, without media supplementation engineered muscles of this type have relatively low numbers of myotubes and produce functional output close to the limit of detection reducing the usefulness of this measure. To further improve the quality of engineered tissues growth factor supplements (FGF2) and commercial supplement cocktails (B-27) were used. B-27 supplement is widely used in primary neuronal and stem cell cultures (Martin *et al*, 2015; Jiwlawat *et al*, 2017). The supplement contains a broad range of components aimed at promoting redox balance, increasing metabolic flexibility, providing trace nutrients and driving cell growth with supplemental growth factors (Insulin) and steroids (triiodothyronine (T3), corticosterone and progesterone). The roles of these components are broad with protection from redox stress and provision of trace nutrients likely to promote cell survival and lead to increased cell numbers (Shaban *et al*, 2017). Growth factors and steroid hormones are more likely to have cell type specific effects, with insulin having been shown in myoblasts to increase fusion and the expression of myogenic genes, as well as driving proliferation (Saini *et al*, 2018; Wu *et al*, 2014). Thyroid hormones have well established roles in regulating skeletal muscle growth and differentiation, with hypothyroidism leading to reduced muscle mass (Bloise *et al*, 2018). At a molecular level the action of T3 on the expression of myogenic genes, MyoD and Myogenin, has been clearly established suggesting T3 will drive myogenesis in engineered muscles (Muscat *et al*, 1994; Downes *et al*, 1993). The effects of progesterone on skeletal muscle remain relatively unexamined, with the expression of progesterone receptors yet to be confirmed in myoblasts (Kim *et al*, 2016). Finally, corticosterone, which has only a limited role in humans with its ortholog cortisol being the predominant corticosteroid, drives skeletal muscle atrophy and inhibits insulin signalling *in vivo* and may perform similar function in culture (Braun & Marks, 2015). The paucity of data makes the action of progesterone and the apparent adversarial effects of insulin and corticosterone make it difficult to unpick precisely the role of each component in B-27 and future work may be required to better understand the mechanisms by which the supplement drives myogenesis. Previous work using B-27 with skeletal muscle myoblasts has been shown to promote myoblast survival, but not differentiation in primary rat MPCs (Das *et al*, 2009) while as a serum replacement for iPSCs B-27 promotes myogenesis (Jiwlawat *et al*, 2017). In the first case B-27 was included throughout culture and may better compare with the addition of B-27 to the growth phase in this work where increases in cell number were observed but no increase in myogenesis. Whereas in the work by Jiwlawat et al, the addition of B-27 occurs to support terminal differentiation, which is similar to the addition of B-27 in the differentiation phase in this work. The growth factor FGF2 is known to promote myoblast proliferation, matrix remodelling and attachment and is widely used in MPC culture (Yablonka-Reuveni & Rivera, 1994; Bakooshli *et al*, 2019) whilst also inhibiting differentiation (Clegg *et al*, 1987; Allen *et al*, 1984) and so was only used in the growth phase of engineered muscles. A doubling in force production shows an effect of supplementation, but this is not driven by an increase in nuclei at maturity. It is possible proliferation may happen earlier in FGF2 supplemented muscles or that there is a priming effect of FGF2 which leads to increased maturity and so force production. The data presented here does not allow this to be examined further. Together this data demonstrates that supplementation effectively improves the quality of engineered muscle (MyHC coverage and force production) produced from hMPCs and although this may not be the absolute optimal supplementation protocol presents a viable solution to produce functional human engineered muscle (Fig 2).

Although myotube CSA and MyHC coverage are low compared to somatic human muscle (Lexell & Taylor, 1991), morphological examination of engineered tissues reveals an organised basement membrane, shown by laminin rings surrounding muscle fibres and the presence of Pax7 positive nuclei within a niche comparable to that seen *in vivo* (Fig 3). However, the niche present in native muscle contains approximately 5-8% of Pax7+ nuclei, substantially greater (10/15-fold) than observed in control engineered tissues (Snow, 1981; Lindström *et al*, 2015). This may be explained by a lack of developmental cues such as the exercise/injury seen *in vivo* which activate satellite cells and cause proliferation (Macaluso *et al*, 2013). This stem cell niche, however rare, is a key feature of skeletal muscle, underpinning the regenerative capacity of skeletal muscle and supporting tissue growth and turnover, and therefore should be present in models of engineered skeletal muscle (Seale *et al*, 2000; Moss & Leblond, 1971; Mauro, 1961).

To examine if these engineered tissues follow similar patterns of regeneration as native skeletal muscle, myogenic and non-myogenic markers were examined through protein and mRNA expression. During the initial 2-days following injury, a proliferative response of MyoD+ myoblasts was observed, before a return to pre-injury levels, a response consistent with *in vivo* data. However, no increase in Pax7+ nuclei was observed during this period as would be expected from *in vivo* data (Tian *et al*, 2016). This lack of Pax7 proliferation could be due to a lack of activation of these cells following injury, with regeneration driven instead by unfused MyoD+ MPCs, or due to the very low percentages of Pax7+ nuclei present making any changes difficult to detect. Distinguishing between these possibilities is difficult. However the increases in mRNA expression of Pax7 and Pax7+ nuclei later in regeneration suggest a capacity to expand this cell population, increase the frequency of the niche and potentially mimic the self-renewing capacity of satellite cells *in vivo* (Kuang *et al*, 2007, 2008).

The self-renewal capacity of Pax7+ cells *in vivo* is highly complex, relying upon asymmetric division of satellite cells, and has not been examined in detail here (Kuang *et al*, 2007, 2008). However, the post injury proportion of Pax7+ nuclei (2.1%) is more closely aligned to *in vivo* proportions (4-7%) than the pre-injury (0.4%), suggesting that an injury stimulus may be required to trigger self-renewal and population expansion. This cannot however be absolutely confirmed without the equivalent of a contralateral control, which is not included. Consequently, we highlight that the increased prevalence of Pax7+ nuclei observed could be due to repeated growth and differentiation phases, and not solely driven by the injury response. The data presented shows that engineered human tissues can mimic some of the key events of regeneration, including the expansion of MPCs and the renewal of the Pax7+ cell population alongside the recovery of function and myotubes (Fig4/5).

Non-myogenic markers Pparg and Runx2 drive non-regenerative repair defects *in vivo* (Lin *et al*, 2011; Dammone *et al*, 2018). Pparg showed an upregulation across regeneration, although this was lower in magnitude than non-regenerative expression *in vivo* (Dammone *et al*, 2018). Runx2 does not show significant upregulation although some variation from control is observed (Fig 5). As remixed engineered muscles contain all cell types obtained from skeletal muscle, these expression patterns may represent the expansion, or increased activity of non-regenerative cells types, which should allow future work to examine how these populations progress to develop non-regenerative defects and how they may be manipulated to improve clinical outcomes.

The model presented here provides a platform to generate large numbers of tissue engineered muscles from a single microbiopsy. Utilising CD56 sorting and media supplementation this protocol is robust and allows researchers, in combination with the open source mould system, to rapidly generate human skeletal muscle tissues within their laboratory (Rimington *et al*, 2018). The demonstration that these tissues regenerate following chemical insult allows the study of human skeletal muscle regeneration, including cell population dynamics across time, to be undertaken without the need to invasively sample patients repeatedly. In addition, the flexibility of the system allows for future work to build complexity, such as through the addition of immune cells to simulate an inflammatory response (Juhas *et al*, 2018a) or mechanical/electrical stimulation to capture the effects of post-injury exercise (Aguilar-Agon *et al*, 2019; Khodabukus *et al*, 2019). As the complexity and maturity of these models develops, they will present an opportunity to test putative clinical interventions in a high throughput manner on human tissue, adding a novel tool to the preclinical testing tool box to help improve lead screening and ultimately improve healthcare for patients.

## Abbreviations

GM: Growth medium
DM: Differentiation medium
ECM: Extracellular matrix
MPC: Myogenic precursor cell
hMPC: Human myogenic precursor cell
MyHC: Myosin heavy chain
SC: Satellite cell
VML: Volumetric muscle loss
FAP: Fibro/adipogenic cell
3D: Three dimensional
PSC: Pluripotent stem cell
DMEM: Dulbecco’s Modified Eagle’s Medium
FBS: Fetal Bovine Serum
P/S: Penicillin/Streptomycin
MEM: Minimal essential medium
PBS: Phosphate buffered saline
KRH: Krebs-Ringer-HEPES
BaCl_2_: Barium Chloride

## Acknowledgements

The authors would like to acknowledge Loughborough University and EPSRC (grant reference EP/N509516/1) for funding and support for this work.

## Author contributions

J.F. contributed to the design of experiments, performed all experimental work and data analysis and drafted the manuscript. A.C., R.R. and O.D. contributed significantly to experimental design and critically reviewed the manuscript throughout the drafting process. P.W. critically reviewed the manuscript and conducted all biopsies for this study. M.L. conceived the concept of the work, contributed to experimental design and critically reviewed the manuscript.

## References

Agarwal S, Cholok D, Loder S, Li J, Breuler C, Chung MT, Sung HH, Ranganathan K, Habbouche J, Drake J, Peterson J, Priest C, Li S, Mishina Y & Levi B (2016) mTOR inhibition and BMP signaling act synergistically to reduce muscle fibrosis and improve myofiber regeneration. JCI Insight 1: 1–12

Agrawal G, Aung A & Varghese S (2017) Lab on a Chip evaluate tissue formation and injury †. Lab Chip

Aguilar-Agon KW, Capel AJ, Martin NRW, Player DJ & Lewis MP (2019) Mechanical loading stimulates hypertrophy in tissue-engineered skeletal muscle: Molecular and phenotypic responses. J. Cell. Physiol. 234: 23547–23558

Allen RE, Dodson M V. & Luiten LS (1984) Regulation of skeletal muscle satellite cell proliferation by bovine pituitary fibroblast growth factor. Exp. Cell Res. 152: 154–160

Bakooshli MA, Lippmann ES, Mulcahy B, Iyer N, Nguyen CT, Tung K, Stewart BA, Van Den Dorpel H, Fuehrmann T, Shoichet M, Bigot A, Pegoraro E, Ahn H, Ginsberg H, Zhen M, Ashton RS & Gilbert PM (2019) A 3d culture model of innervated human skeletal muscle enables studies of the adult neuromuscular junction. Elife 8: 1–29

Belmont PJ, McCriskin BJ, Sieg RN, Burks R & Schoenfeld AJ (2012) Combat wounds in Iraq and Afghanistan from 2005 to 2009. J. Trauma Acute Care Surg. 73: 3–12

Bi P, Ramirez-Martinez A, Li H, Cannavino J, McAnally JR, Shelton JM, Sánchez-Ortiz E, Bassel-Duby R & Olson EN (2017) Control of muscle formation by the fusogenic micropeptide myomixer. Science (80-.). 356: 323–327

Bloise FF, Cordeiro A & Ortiga-Carvalho TM (2018) Role of thyroid hormone in skeletal muscle physiology. J. Endocrinol. 236: R57–R68

Brady MA, Lewis MP & Mudera V (2008) Synergy between myogenic and non-myogenic cells in a 3D tissue-engineered craniofacial skeletal muscle construct. J. Tissue Eng. Regen. Med. 2: 408–17

Braun TP & Marks DL (2015) The regulation of muscle mass by endogenous glucocorticoids. Front. Physiol. 6: 1–12

Capel AJ, Rimington RP, Fleming JW, Player DJ, Baker LA, Turner MC, Jones JM, Martin NRW, Ferguson RA, Mudera VC & Lewis MP (2019) Scalable 3D Printed Molds for Human Tissue Engineered Skeletal Muscle. Front. Bioeng. Biotechnol. 7: 20

Clegg CH, Linkhart TA, Olwin BB & Hauschka SD (1987) Growth factor control of skeletal muscle differentiation: Commitment to terminal differentiation occurs in G1 phase and is repressed by fibroblast growth factor. J. Cell Biol. 105: 949–956

Dammone G, Karaz S, Lukjanenko L, Winkler C, Sizzano F, Jacot G, Migliavacca E, Palini A, Desvergne B, Gilardi F & Feige JN (2018) PPARγ controls ectopic adipogenesis and cross-talks with myogenesis during skeletal muscle regeneration. Int. J. Mol. Sci. 19: 1–17

Das M, Rumsey JW, Bhargava N, Gregory C, Riedel L, Kang JF & Hickman JJ (2009) Developing a novel serum-free cell culture model of skeletal muscle differentiation by systematically studying the role of different growth factors in myotube formation. In Vitro Cell. Dev. Biol. Anim. 45: 378–387

Dharm-Datta S & McLenaghan J (2013) Medical lessons learnt from the US and Canadian experience of treating combat casualties from Afghanistan and Iraq. J. R. Army Med. Corps 159: 102–9

Downes M, Griggs R, Atkins A, Olson EN & Muscat GEO (1993) Identification of a thyroid hormone response element in the mouse myogenin gene: Characterization of the thyroid hormone and retinoid X receptor heterodimeric binding site. Cell Growth Differ. 4: 901–910

Dumont NA, Wang YX & Rudnicki MA (2015) Intrinsic and extrinsic mechanisms regulating satellite cell function. Dev. 142: 1572–1581

EU (2010) Directive 2010/63/EU of the European Parliament and of the Council of 22 September 2010 on the protection of animals used for scientific purposes. Off. J. Eur. Union L 276, 20.:

Fiore D, Judson RN, Low M, Lee S, Zhang E, Hopkins C, Xu P, Lenzi A, Rossi FM V & Lemos DR (2016) Pharmacological blockage of fi bro / adipogenic progenitor expansion and suppression of regenerative fi brogenesis is associated with impaired skeletal muscle regeneration. Stem Cell Res. 17: 161–169

Fleming JW, Capel AJ, Rimington RP, Player DJ, Stolzing A & Lewis MP (2019) Functional regeneration of tissue engineered skeletal muscle in vitro is dependent on the inclusion of basement membrane proteins. Cytoskeleton 76: 371–382

Fournier-Farley C, Lamontagne M, Gendron P & Gagnon DH (2016) Determinants of Return to Play after the Nonoperative Management of Hamstring Injuries in Athletes. Am. J. Sports Med. 44: 2166–2172

Freckleton G & Pizzari T (2013) Risk factors for hamstring muscle strain injury in sport: A systematic review and meta-analysis. Br. J. Sports Med. 47: 351–358

Hardy D, Besnard A, Latil M, Jouvion G, Briand D, Thépenier C, Pascal Q, Guguin A, Gayraud-Morel B, Cavaillon J-M, Tajbakhsh S, Rocheteau P & Chrétien F (2016) Comparative Study of Injury Models for Studying Muscle Regeneration in Mice. PLoS One 11: e0147198

Hayot M, Michaud A, Koechlin C, Caron MA, LeBlanc P, Préfaut C & Maltais F (2005) Skeletal muscle microbiopsy: A validation study of a minimally invasive technique. Eur. Respir. J. 25: 431–440

Hindi SM & Kumar A (2016) TRAF6 regulates satellite stem cell self-renewal and function during regenerative myogenesis. J. Clin. Invest. 126: 151–168

Huang Y-C, Dennis RG, Larkin L & Baar K (2005) Rapid formation of functional muscle in vitro using fibrin gels. J. Appl. Physiol. 98: 706–13

Illa I, Leon-Monzon M & Dalakas MC (1992) Regenerating and denervated human muscle fibers and satellite cells express neural cell adhesion molecule recognized by monoclonal antibodies to natural killer cells. Ann. Neurol. 31: 46–52

Järvinen TA, Järvinen M & Kalimo H (2013) Regeneration of injured skeletal muscle after the injury. Muscles. Ligaments Tendons J. 3: 337–45

Jiwlawat S, Lynch E, Glaser J, Smit-Oistad I, Jeffrey J, Van Dyke JM & Suzuki M (2017) Differentiation and sarcomere formation in skeletal myocytes directly prepared from human induced pluripotent stem cells using a sphere-based culture. Differentiation 96: 70–81

Joe AW, Yi L, Natarajan A, Le Grand F, So L, Wang J, Rudnicki MA & Rossi FM (2010) Muscle injury activates resident fibro/adipogenic progenitors that facilitate myogenesis. Nat Cell Biol 12: 153–163

Juhas M, Abutaleb N, Wang JT, Ye J, Shaikh Z, Sriworarat C, Qian Y & Bursac N (2018a) Incorporation of macrophages into engineered skeletal muscle enables enhanced muscle regeneration. Nat. Biomed. Eng. 2: 942–954

Juhas M, Abutaleb N, Wang JT, Ye J, Shaikh Z, Sriworarat C, Qian Y & Bursac N (2018b) Incorporation of macrophages into engineered skeletal muscle enables enhanced muscle regeneration. Nat. Biomed. Eng.: 1

Juhas M, Engelmayr GCJ, Fontanella a. N, Palmer GM & Bursac N (2014) Biomimetic engineered muscle with capacity for vascular integration and functional maturation in vivo. Proc Natl Acad Sci U S A 111: 5508–13

Khodabukus A, Madden L, Prabhu NK, Koves TR, Jackman CP, Muoio DM & Bursac N (2019) Electrical stimulation increases hypertrophy and metabolic flux in tissue-engineered human skeletal muscle. Biomaterials 198: 259–269

Kim YJ, Tamadon A, Park HT, Kim H & Ku S-Y (2016) The role of sex steroid hormones in the pathophysiology and treatment of sarcopenia. Osteoporos. Sarcopenia 2: 140–155

Kuang S, Gillespie MA & Rudnicki MA (2008) Niche Regulation of Muscle Satellite Cell Self-Renewal and Differentiation. Cell Stem Cell 2: 22–31

Kuang S, Kuroda K, Le Grand F & Rudnicki MA (2007) Asymmetric Self-Renewal and Commitment of Satellite Stem Cells in Muscle. Cell 129: 999–1010

Lemos DR, Babaeijandaghi F, Low M, Chang C-K, Lee ST, Fiore D, Zhang R-H, Natarajan A, Nedospasov S a & Rossi FM V (2015) Nilotinib reduces muscle fibrosis in chronic muscle injury by promoting TNF-mediated apoptosis of fibro/adipogenic progenitors. Nat. Med. 21: 786–794

Lepper C, Partridge TA & Fan CM (2011) An absolute requirement for pax7-positive satellite cells in acute injury-induced skeletal muscle regeneration. Development 138: 3639–3646

Lexell J & Taylor CC (1991) Variability in muscle fibre areas in whole human quadriceps muscle: effects of increasing age. J. Anat. 174: 239–49

Lin L, Shen Q, Leng H, Duan X, Fu X & Yu C (2011) Synergistic inhibition of endochondral bone formation by silencing Hif1α and Runx2 in trauma-induced heterotopic ossification. Mol. Ther. 19: 1426–1432

Lindström M, Tjust AE & Domellöf FP (2015) Pax7-positive cells/satellite cells in human extraocular muscles. Investig. Ophthalmol. Vis. Sci. 56: 6132–6143

Macaluso F, Brooks NE, Niesler CU & Myburgh KH (2013) Satellite Cell Pool Expansion Is Affected By Skeletal Muscle Characteristics. Muscle and Nerve 48: 109–116

Madden L, Juhas M, Kraus WE, Truskey GA & Bursac N (2015) Bioengineered human myobundles mimic clinical responses of skeletal muscle to drugs. Elife 4: e04885

Maffioletti SM, Sarcar S, Henderson ABH, Mannhardt I, Pinton L, Moyle LA, Steele-Stallard H, Cappellari O, Wells KE, Ferrari G, Mitchell JS, Tyzack GE, Kotiadis VN, Khedr M, Ragazzi M, Wang W, Duchen MR, Patani R, Zammit PS, Wells DJ, et al (2018) Three-Dimensional Human iPSC-Derived Artificial Skeletal Muscles Model Muscular Dystrophies and Enable Multilineage Tissue Engineering. Cell Rep. 23: 899–908

Mann CJ, Perdiguero E, Kharraz Y, Aguilar S, Pessina P, Serrano AL & Muñoz-Cánoves P (2011) Aberrant repair and fibrosis development in skeletal muscle. Skelet. Muscle 1: 21

Martin NRW, Passey SL, Player DJ, Mudera V, Baar K, Greensmith L & Lewis MP (2015) Neuromuscular Junction Formation in Tissue-Engineered Skeletal Muscle Augments Contractile Function and Improves Cytoskeletal Organization. Tissue Eng. Part A 21: 2595–2604

Mauro A (1961) Satellite cell of skeletal muscle fibers. J. Biophys. Biochem. Cytol. 9: 493–5

Merly F, Lescaudron L, Rouaud T, Crossin F & Gardahaut MF (1999) Macrophages enhance muscle satellite cell proliferation and delay their differentiation. Muscle and Nerve 22: 724–732

Millay DP, O’Rourke JR, Sutherland LB, Bezprozvannaya S, Shelton JM, Bassel-Duby R & Olson EN (2013) Myomaker is a membrane activator of myoblast fusion and muscle formation. Nature 499: 301–305

Morimoto Y, Onoe H & Takeuchi S (2018) Biohybrid robot powered by an antagonistic pair of skeletal muscle tissues. Sci. Robot. 3: eaat4440

Moss FP & Leblond CP (1971) Satellite cells as the source of nuclei in muscles of growing rats. Anat. Rec. 170: 421–35

Mueller AA, Van Velthoven CT, Fukumoto KD, Cheung TH & Rando TA (2016) Intronic polyadenylation of PDGFRα in resident stem cells attenuates muscle fibrosis. Nat. Publ. Gr. 540: 276–279

Muscat GEO, Mynett-johnson L, Dowhan D, Downes M & Griggs R (1994) Activation of myoD gene transcription by 3,5,3’-triiodo-L-thyronine: A direct role for the thyroid hormone and retinoid X receptors. Nucleic Acids Res. 22: 583–591

Olson LE & Soriano P (2009) Increased PDGFRalpha activation disrupts connective tissue development and drives systemic fibrosis. Dev. Cell 16: 303–13

Quinn ME, Goh Q, Kurosaka M, Gamage DG, Petrany MJ, Prasad V & Millay DP (2017) Myomerger induces fusion of non-fusogenic cells and is required for skeletal muscle development. Nat. Commun. 8: 1–9

Rao L, Qian Y, Khodabukus A, Ribar T & Bursac N (2018) Engineering human pluripotent stem cells into a functional skeletal muscle tissue. Nat. Commun. 9: 1–12

Rimington R, Capel AJ, Fleming J, Player D, Mudera V, Jones J, Martin NRW, Turner M, Baker L, Ferguson R & Lewis M (2018) 50uL FDM removable insert.

Rimington RP, Capel AJ, Christie SDR & Lewis MP (2017) Biocompatible 3D printed polymers: Via fused deposition modelling direct C2C12 cellular phenotype in vitro. Lab Chip 17: 2982–2993

Rose PE (2002) Pathology of Skeletal Muscle, 2nd ed: Carpenter S, Karpati G. (£140.00.) Oxford University Press, 2001. ISBN 0 19 506364 3. J. Clin. Pathol. 55: 480

Russell WMS, Burch RL & Hume CW (1959) The principles of humane experimental technique.

Saini A, Rullman E, Lilja M, Mandić M, Melin M, Olsson K & Gustafsson T (2018) Asymmetric cellular responses in primary human myoblasts using sera of different origin and specification. PLoS One 13: 1–16

Schindelin J, Arganda-Carreras I, Frise E, Kaynig V, Longair M, Pietzsch T, Preibisch S, Rueden C, Saalfeld S, Schmid B, Tinevez J-Y, White DJ, Hartenstein V, Eliceiri K, Tomancak P & Cardona A (2012) Fiji: an open-source platform for biological-image analysis. Nat Meth 9: 676–682

Schmittgen TD & Livak KJ (2008) Analyzing real-time PCR data by the comparative CT method. Nat. Protoc. 3: 1101–1108

Seale P, Sabourin LA, Girgis-Gabardo A, Mansouri A, Gruss P & Rudnicki MA (2000) Pax7 is required for the specification of myogenic satellite cells. Cell 102: 777–86

Segawa M, Fukada S ichiro, Yamamoto Y, Yahagi H, Kanematsu M, Sato M, Ito T, Uezumi A, Hayashi S, Miyagoe-Suzuki Y, Takeda S, Tsujikawa K & Yamamoto H (2008) Suppression of macrophage functions impairs skeletal muscle regeneration with severe fibrosis. Exp. Cell Res. 314: 3232–3244

Shaban S, El-Husseny MWA, Abushouk AI, Salem AMA, Mamdouh M & Abdel-Daim MM (2017) Effects of Antioxidant Supplements on the Survival and Differentiation of Stem Cells. Oxid. Med. Cell. Longev. 2017:

Snow MH (1981) Satellite cell distribution within the soleus muscle of the adult mouse. Anat. Rec. 201: 463–9

Summan M, Warren GL, Mercer RR, Chapman R, Hulderman T, Van Rooijen N & Simeonova PP (2006) Macrophages and skeletal muscle regeneration: a clodronate-containing liposome depletion study. Am J Physiol Regul Integr Comp Physiol 290: R1488–95

Tian ZL, Jiang SK, Zhang M, Wang M, Li JY, Zhao R, Wang LL, Li SS, Liu M, Zhang MZ & Guan DW (2016) Detection of satellite cells during skeletal muscle wound healing in rats: time-dependent expressions of Pax7 and MyoD in relation to wound age. Int. J. Legal Med. 130: 163–172

Tiburcy M, Markov A, Kraemer LK, Christalla P, Rave-Fraenk M, Fischer HJ, Reichardt HM & Zimmermann W (2019) Regeneration competent satellite cell niches in rat engineered skeletal muscle. FASEB BioAdvances 1: 731–746

Uezumi A, Fukada S, Yamamoto N, Takeda S & Tsuchida K (2010) Mesenchymal progenitors distinct from satellite cells contribute to ectopic fat cell formation in skeletal muscle. Nat Cell Biol 12: 143–152

Uezumi A, Ito T, Morikawa D, Shimizu N, Yoneda T, Segawa M, Yamaguchi M, Ogawa R, Matev MM, Miyagoe-Suzuki Y, Takeda S, Tsujikawa K, Tsuchida K, Yamamoto H & Fukada S (2011) Fibrosis and adipogenesis originate from a common mesenchymal progenitor in skeletal muscle. J. Cell Sci. 124: 3654–64

Vandenburgh HH, Karlisch P & Farr L (1988) Maintenance of highly contractile tissue-cultured avian skeletal myotubes in collagen gel. In Vitro Cell. Dev. Biol. 24: 166–74

Wangensteen A, Tol JL, Witvrouw E, Van Linschoten R, Almusa E, Hamilton B & Bahr R (2016) Hamstring Reinjuries Occur at the Same Location and Early after Return to Sport. Am. J. Sports Med. 44: 2112–2121

Wheatley BM, Hanley MG, Wong VW, Sabino JM, Potter BK, Tintle SM, Fleming ME & Valerio IL (2015) Heterotopic Ossification following Tissue Transfer for Combat-Casualty Complex Periarticular Injuries. Plast. Reconstr. Surg. 136: 808e–14e

Wosczyna MN, Konishi CT, Perez Carbajal EE, Wang TT, Walsh RA, Gan Q, Wagner MW & Rando TA (2019) Mesenchymal Stromal Cells Are Required for Regeneration and Homeostatic Maintenance of Skeletal Muscle. Cell Rep. 27: 2029–2035.e5

Wu YJ, Fang YH, Chi HC, Chang LC, Chung SY, Huang WC, Wang XW, Lee KW & Chen SL (2014) Insulin and LiCl synergistically rescue myogenic differentiation of FoxO1 over-expressed myoblasts. PLoS One 9:

Xu B, Zhang M, Perlingeiro RCR & Shen W (2019) Skeletal Muscle Constructs Engineered from Human Embryonic Stem Cell Derived Myogenic Progenitors Exhibit Enhanced Contractile Forces When Differentiated in a Medium Containing EGM-2 Supplements. Adv. Biosyst. 1900005: 1–11

Yablonka-Reuveni Z & Rivera a J (1994) Temporal expression of regulatory and structural muscle proteins during myogenesis of satellite cells on isolated adult rat fibers. Dev. Biol. 164: 588–603

Zammit PS (2017) Function of the myogenic regulatory factors Myf5, MyoD, Myogenin and MRF4 in skeletal muscle, satellite cells and regenerative myogenesis. Semin. Cell Dev. Biol. 72: 19–32

