## Supplementary figures for "Bioengineered human skeletal muscle with a Pax7+ satellite cell niche capable of functional regeneration"

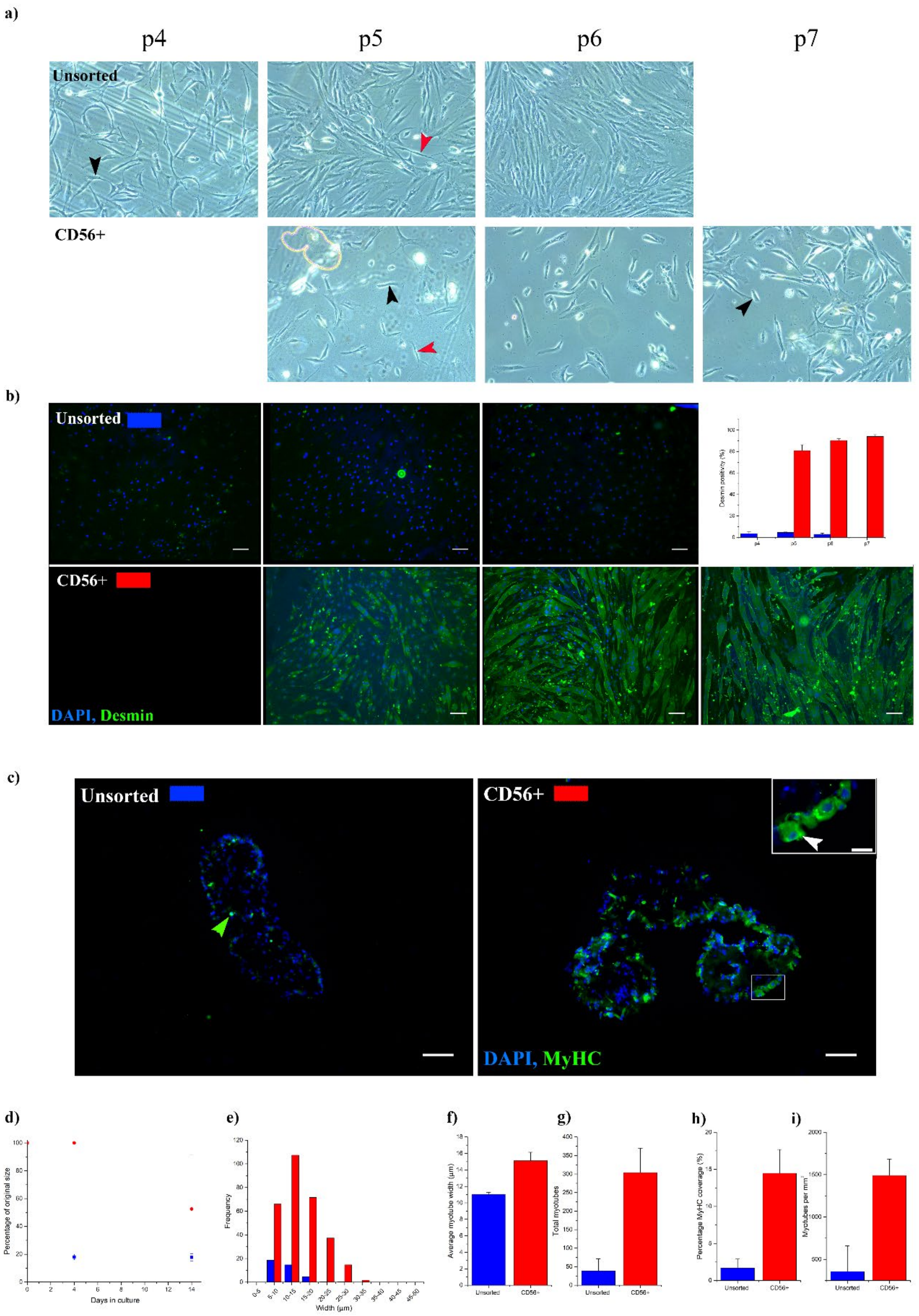

**Table S1: Primer sequence table**

| Gene | NCBI accession no. | Sequence |
|---|---|---|
| POLR2B | NM_000938.3 | Fw: AAGGCTTGGTTAGACAACAG |
| | | Rv: TATCGTGGCGGTTCTTCA |
| PAX7 | NM_013945.3 | Fw: ACCCCTGCCTAACCACATC |
| | | Rv: GCGGCAAAGAATCTTGGAGAC |
| MYOG | NM_002479.6 | Fw: CAGCTCCCTCAACCAGGAG |
| | | Rv: GCTGTGAGAGCTGCATTCG |
| PPARG | NM_015869.5 | Fw: AAAGACAACGGACAAATCAC |
| | | Rv: GGGATATTTTTGGCATACTCTG |
| RUNX2 | NM_001024630.4 | Fw: GCAGTATTTACAACAGAGGG |
| | | Rv: TCCCAAAAGAAGTTTTGCTG |

**Figure S1: CD56 enrichment improves desmin positivity and morphological appearance of tissue engineered constructs. (a)** Phase contrast images of human myogenic precursor cells. Black arrows highlight cells with sort square morphology, red arrows show cells with extended fibroblast like morphology. **(b)** Fluorescent micrographs (10x) of unsorted and CD56 enriched cultures following 6 days in differentiation media. Cultures are stained for desmin (green) and nuclei (blue), scale bar 100μm. Inset graph – desmin positivity (%). **(c)** Fluorescent micrographs (10x) of tissue engineered cross sections. MyHC (green) and nuclei (DAPI) are used to identify myotubes. Scale bar 100μm. Inset shows peripherally nucleated fibre (white arrow), scale bar 25μm.**(d)** Hydrogel deformation expressed as a percentage of size at day 0. **(e)** Average frequency of myotubes of given width within a single cross section of tissue engineered muscle. **(f)** Average myotube width **(g)** Total myotubes per cross section **(h)** Percentage of total cross sectional area occupied my MytHC structures **(i)** Myotube density expressed as myotubes per $mm^2$
